# On an Analogy Between Ephaptic Coupling in Large Tracts and Cross-Modal Integration During Illusory Percept

**DOI:** 10.1101/2022.10.27.509121

**Authors:** Aman Chawla, Salvatore Domenic Morgera

## Abstract

In this paper, the authors report simulations analogous to cross-modal integration which can be modeled by thinking of large (in terms of numbers of axons) axon tracts with distinctly demarcated subsets of axons. In such a tract, when the demarcated regions are separately stimulated they fire in response to the stimulation. Due to extracellular current-based coupling, at a later time, unstimulated axons also begin to cross-talk. The simulation graphical result, though on a faster time scale, is analogous to the multi-channel MEG recordings of “AV-(A+V)” obtained by Shams et. al. When the coupling parameter is weakened, this cross-talk disappears. In Shams et. al., the appearance and disappearance of cross-talk are related to the perception of an illusory flash by the subject. Our extrapolated conclusion is that ephaptic coupling might be a part of the mechanism for cross-talk between brain regions mediating, say, audio and visual-processing. We also study noisy ephaptic coupling and additionally do a comparative study with the Kuramoto model, which yields insight into the exact and detailed mechanism of ephaptic synchronization.

## 1 Introduction

In this section we will point the reader to the psychophysical background of cross-modal integration. Cross-modal integration [1] is the synthesis of inputs from auditory and visual modalities, say, into a unified percept in the cortex. An excellent review article which stretches all the way from 1958 to 2010 is [2]. The sound-induced double-flash illusion was used in [3] to investigate this integration and its impact on visual perception. In the study of the illusion, a subject is presented with a flash (V) and a tone (A) separately, and simultaneously (AV). By comparing the results in these different cases, the authors demonstrate that the modalities influence one another and they present physiological recordings which provide evidence for this influence.

There are a number of other additional studies on cross-modal integration, especially those involving the auditory and visual modalities, such as [4]. Most of these studies approach the investigation from the experimental perspective, without developing mathematical models that explain the observations. In the present paper we report that ephaptic coupling in axon tracts that are sub-divided into different regions can be viewed as a prototypical basis for cross-talk between these regions. From a computational angle, the work extends the work first begun in [5] where the authors presented a new geometric model for ephaptic or current-mediated coupling. Further, the work is not new in suggesting that synchronization between neurons has a functional role. See, for example, [6] and the references therein.

The paper is organized as follows. In Section 2 we mimic Figures 2 and 4 of [3] by using a tract of N=30 coupled axons. In Section 3 we add noise to the mix and find that there is long-term dissipation of coupling in the presence of noise. In Section 4 we do a comparative study of [5] and the Kuramoto model, using this study to guide our simulation-based investigation of cross-modal coupling. Finally, we present the limitations of our work and conclude in Section 5. An Appendix reproduces two of the cited figures.

**Figure 1:**
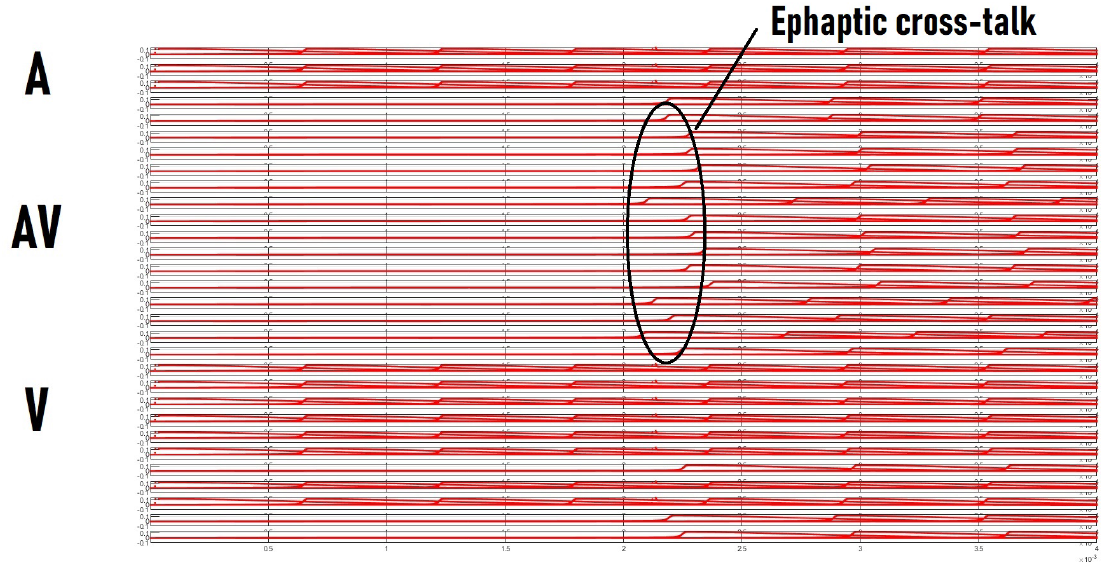
N=30 axons coupled with a strong ratio. Simultaneous stimulation of A and V leads to a delayed response in AV. Compare Figure 2 in [3]. The x-axes represent time in seconds and the y-axes of the various sub-plots represent voltage in Volts.

**Figure 2:**
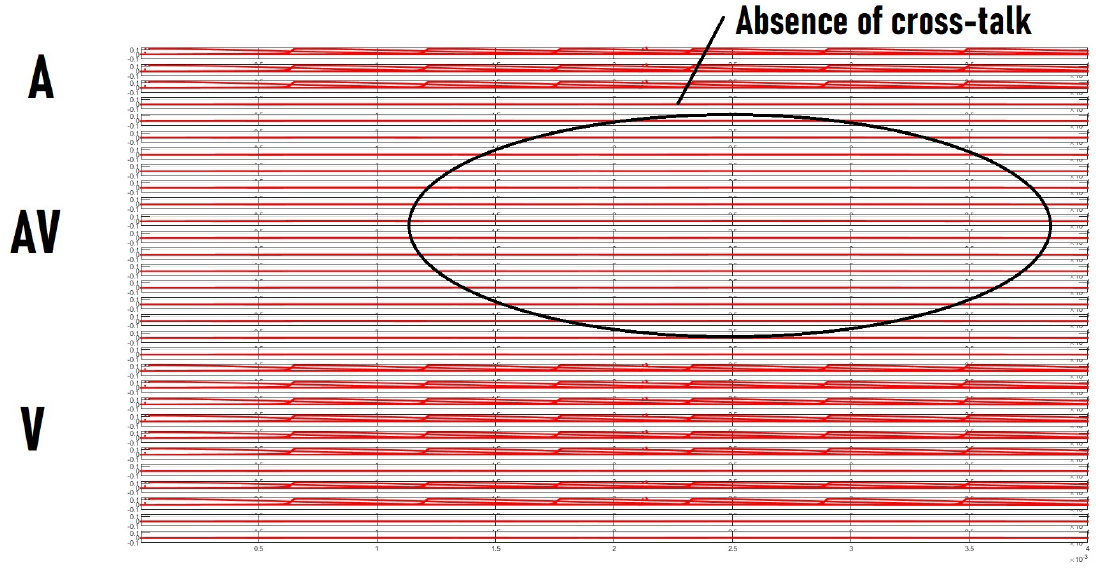
N=30 axons coupled with a weak ratio. Simultaneous stimulation of A and V does not lead to any response in AV. Compare Figure 4 in [3]. The x-axes represent time in seconds and the y-axes of the various sub-plots represent voltage in Volts.

**Figure 3:**
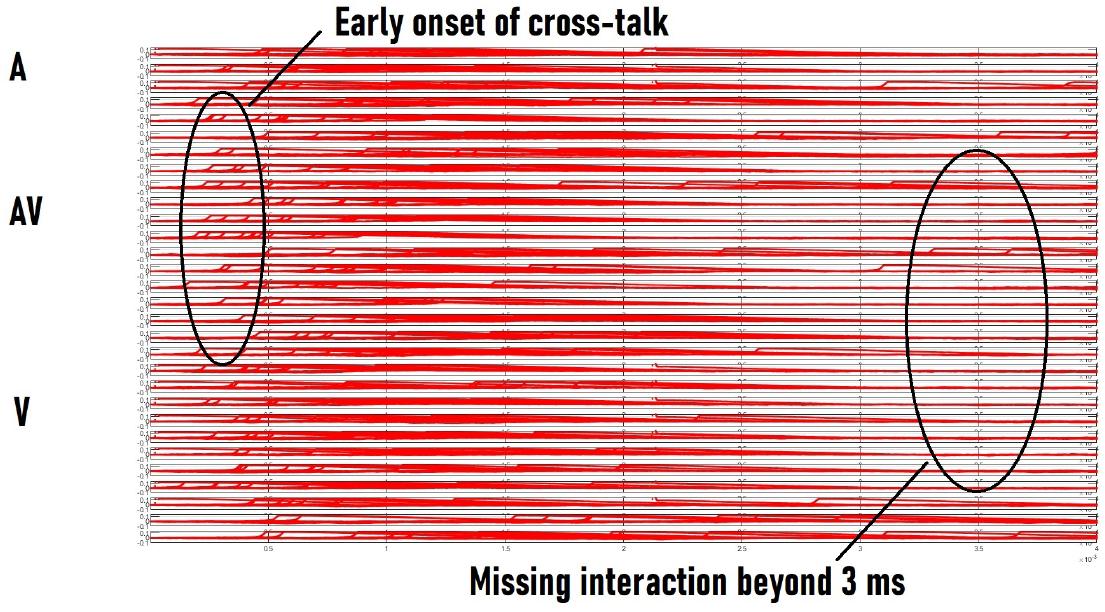
30 axons coupled with a scale of noise = .1e-5 and intermediate ratio = 0.52. The top stimulated axons can be considered to be the A region and the bottom stimulated axons as the V region, with AV being the intermediate unstimulated axons. The x-axes represent time in seconds and the y-axes of the various sub-plots represent voltage in Volts.

**Figure 4:**
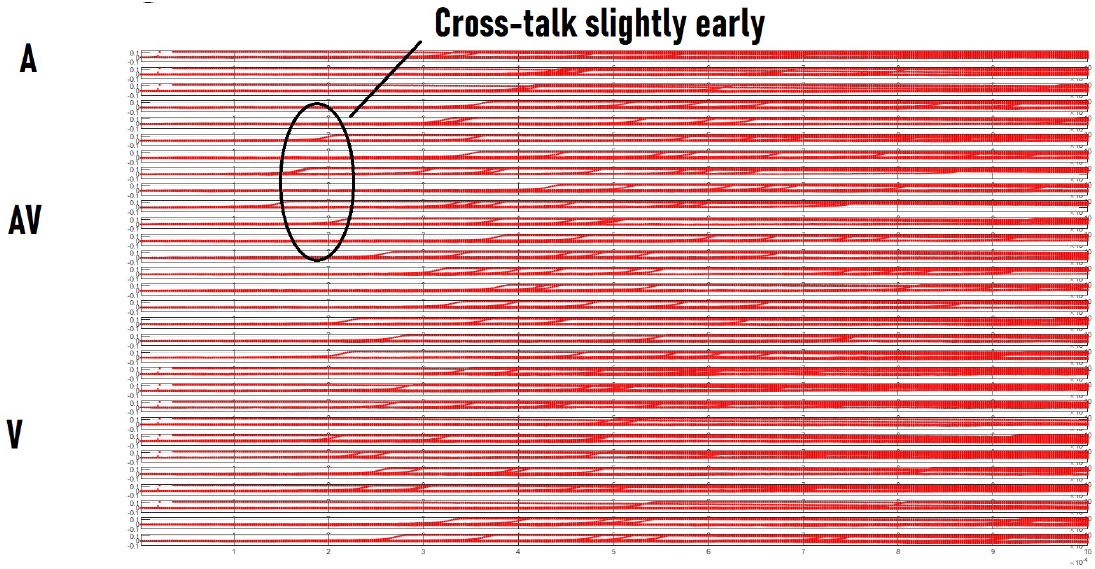
30 axons interacting at a low coupling (ratio = 0.32). The top stimulated axons can be considered to be the A region and the bottom stimulated axons as the V region, with AV being the intermediate unstimulated axons. The x-axes represent time in seconds and the y-axes of the various sub-plots represent voltage in Volts.

### Notation

The notation used in this paper is presented succinctly in Table 1.

**Table 1:** Table of Notation

| S.No. | Symbol | Meaning |
| --- | --- | --- |
| 1 | $c_i$ | Capacitance per unit length of the $i$-th axon |
| 2 | $V_i$ | Transmembrane voltage of the $i$-th axon |
| 3 | $(t, z)$ | Temporal and spatial coordinates |
| 4 | $G_i^{ax}, G_i^{my}$ | (Axoplasmic, myelin-sheath) conductance per unit length for the $i$-th axon |
| 5 | $\theta_i$ | Inclination of the $i$-th axon with respect to the tract axis |
| 6 | $i, p$ | Index of axons, taking values in $\{1, 2, \dots, N\}$ where $N$ is the total number of axons in the tract |
| 7 | $W_{ip}$ | $(i, p)$-th entry of a matrix which is based on the perpendicular distance between axon $i$ and axon $p$ |
| 8 | $K_p$ | Term for the $p$-th axon, defined in [5], based on its inclination and conductance |
| 9 | $I_i^{inj}$ | Injection and ionic current for the $i$-th axon's nodal equation |

## 2 Ephaptic Coupling as a Mechanism for Cross-modal Integration

Ephaptic coupling is a phenomenon within axon tracts where the propagation on different axons becomes synchronized due to coupling extracellular currents [7]. The functional role of this coupling remains a point of debate [8]. In [9] it was proposed that it serves to amplify entropy of the input stimulus to the tract. While entropy is being amplified on the information theoretic or data layer level, what is happening at the physical layer level? The present paper delves into this question.

Figures 1 and 2 suggest that an axon tract can be used as a good test bed to replicate the performance of larger regions of the brain when studied for auditory-visual cross-modal integration. The model used in these figures was first introduced in [5]. In both these figures, the axons are numbered 1 through 30 from the topmost subplot to the bottom-most subplot. Our study is aided by labeling the top few stimulated axons as (A), the bottom few as (V) and the middle unstimulated ones as (AV), even though all belong to the same tract and true adjacency is captured by the W-matrix. The x-axis is time in seconds and the y-axes are voltage in Volts. The range of the x-axis is from 0 to 4 ms.

This should be contrasted with the temporal range in Figure 2 and Figure 4 of Shams et. al. [3]. There the range goes up to 400 ms. If we scale up our figures’ x-axes a hundredfold, the features of interest (cross-talk timings) are more or less preserved.

## 3 Ephaptic Coupling in the Presence of Noise

In this section we study what happens when low-grade noise is added to a A-V-AV demarcated tract (as in the previous section) in which there is current-based ephaptic coupling. Figure 3 suggests that when a noise scaling factor (a constant pre-multiplying the standard normal random variable) of 1e-6 is applied to the axons in a tract at the ion channel level, long-term ephaptic coupling based cross-talk between axons gets suppressed. If we compare this figure with Figure 1, we can see the “missing” interaction beyond 3 ms post-stimulus. It should be reiterated here that the time scales are about 100 fold shorter compared with those in [3].

As a note, in the quantum regime, noise is sometimes seen as useful in conducting information [10]. It would be interesting to see in this context, if the ion channel noise in an axon tract can be viewed as beneficial rather than pernicious. This would involve a capacity or information-theoretic investigation and as such is beyond the scope of the present paper.

## 4 Mathematical Analysis of Ephaptic Synchronization

The analysis in the previous sections has been guided vaguely by a notion of coupling ratio strength or coupling parameter. The ambiguity around this notion must be clarified. In this section, guided by the presentation in [11] and [12], we try to understand ephaptic synchronization quantitatively. Towards this end, consider the following development.

From [5] we note down the governing equation for ephaptic interaction

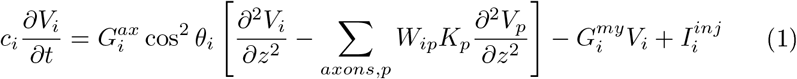

where the notation is explained in Table 1. We juxtapose Equation (1) with the following equation from [11]:

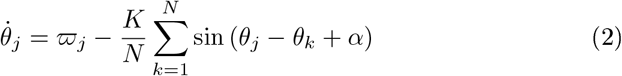

for *j* = 1, …, *N*, where *θ*_*j*_ is the phase of the *j*-th oscillator, *ϖ*_*j*_ is its bare frequency, *K* is the coupling constant, and *α* is a constant. This suggests to us to make the following identification,

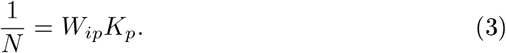

The identification above leads Equation (1) into the form

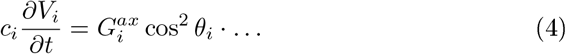

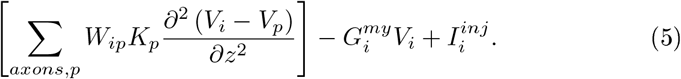

Thus, under identification (3), we have obtained a Kuramoto-like oscillator equation from the geometric ephaptic equation. We are thus aided in our mathematical understanding of ephaptic coupling in the *Kuramoto regime*. Using Equation (3) again, we rewrite Equation (5) as

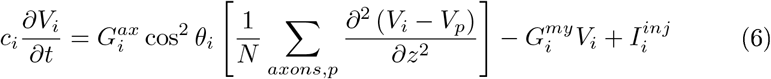

We next assume that the myelin conductance is negligible compared to the axoplasmic conductance. This enables us to write

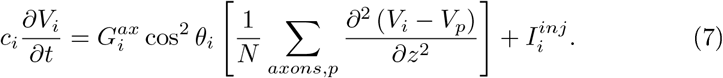

From this last equation we come to the conclusion that 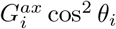 plays the role of the coupling constant *K* in the Kuramoto equation, Equation (2). Thus the axoplasmic resistivity, which is proportional to the parameter ‘ratio’ in our program [5], can be varied and we would expect to see changes in the axon-axon coupling and synchronization. For low coupling, we have Figure 4 and for high coupling we have Figure 5. In both figures, the x-axes represent time in seconds and the y-axes represent voltage in Volts. The axons are arranged from top to bottom in sub-plots.

**Figure 5:**
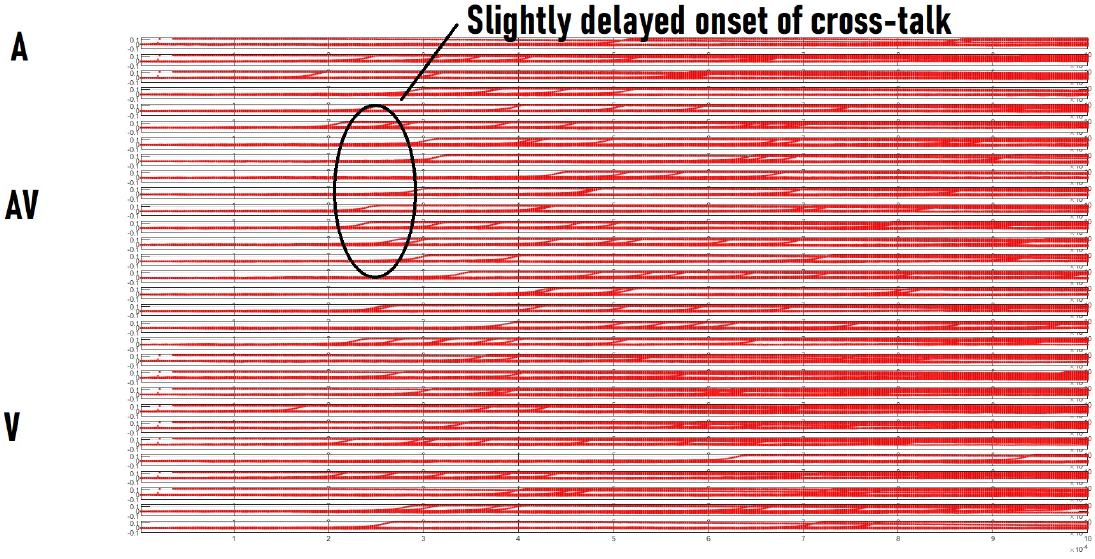
30 axons interacting at a high coupling (ratio = 0.92). The top stimulated axons can be considered to be the A region and the bottom stimulated axons as the V region, with AV being the intermediate unstimulated axons. Paradoxically, onset of cross-talk appears delayed compared to the low coupling case. The x-axes represent time in seconds and the y-axes of the various sub-plots represent voltage in Volts.

## 5 Conclusion

Ephaptic coupling can be used to shed light on the underlying mechanism of cross-modal integration. It is suggested that extracellular currents or electric fields lead to coupling between different brain cortices, coupling which ultimately binds the modalities represented in these cortices. This imparts a new and valuable role to these extracellular currents and fields.

To re-emphasize, while there has been past work on cross-modal integration from the psychophysics-experimental perspective, this is the first paper, to our knowledge, which proposes that current-mediated-coupling in the geometric context can be seen as a potential mechanism by which axons, and by extrapolation (to fields), brain regions, tend to begin to talk to one another, and coordinate their activity within a certain temporal latency. The latency is dependent on the geometry of the problem and the nature or strength of the implemented coupling.

There are several limitations to our work. We didn’t study larger-sized axon tracts; this can be overcome in future work. Furthermore, we might be able to replicate the geometry of the actual regions in the human brain where integration takes place, to follow that shown by experiments, and this can be investigated in future work. Specifically, we would like to see which geometry brings the latency up to 400 milliseconds or so. In future work, we might also build a more general mathematical model for cross-modal integration than the one suggested here, by modeling different brain regions as *diferent* axon tracts, and allowing those tracts to be coupled via electric fields. Another limitation of the present paper is a true statistical analysis, based on noisy simulations, where we actually compute the p-values of “AV-(A+V)” as is done in the psychophysics literature. This can be addressed in future work.

In summary, this work has expanded our understanding of the function of ephaptic coupling in the brain. Forms of coupling such as the one studied here are shown to be potentially analogous to the inter-regional processing that obtains during ‘binding’ or integration of modalities, as shown in the Appendix.

## Appendix

### AV-(A+V) during illusion

**Figure 6:**
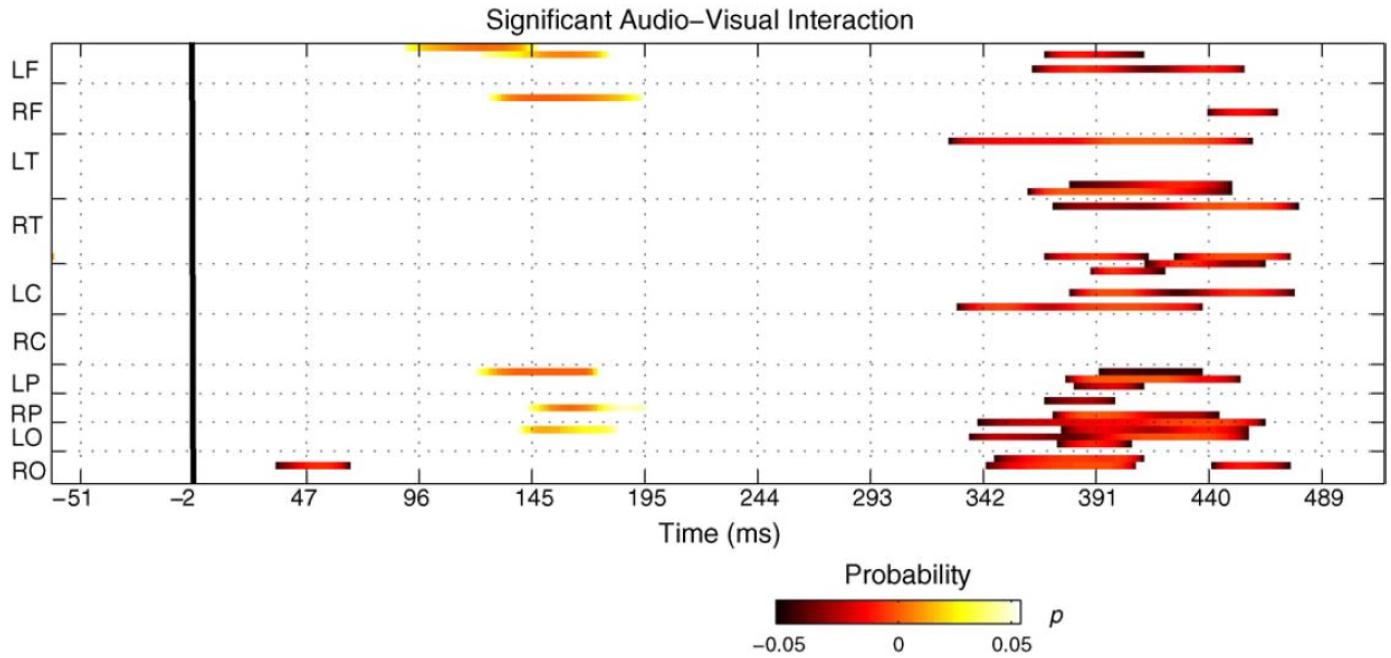
Significant crossmodal interactions obtained by displaying p-values for AV-(A+V), for the case when the illusion was perceived. Reproduced, with permission, from [3]. Whereas only the occipital and temporal lobes are directly connected to the sensory modalities of vision and audition, when the illusion is perceived, the whole brain (all regions) shows activity around 400 milliseconds. Compare with the figures in the main text.

### AV-(A+V) in non-illusion

**Figure 7:**
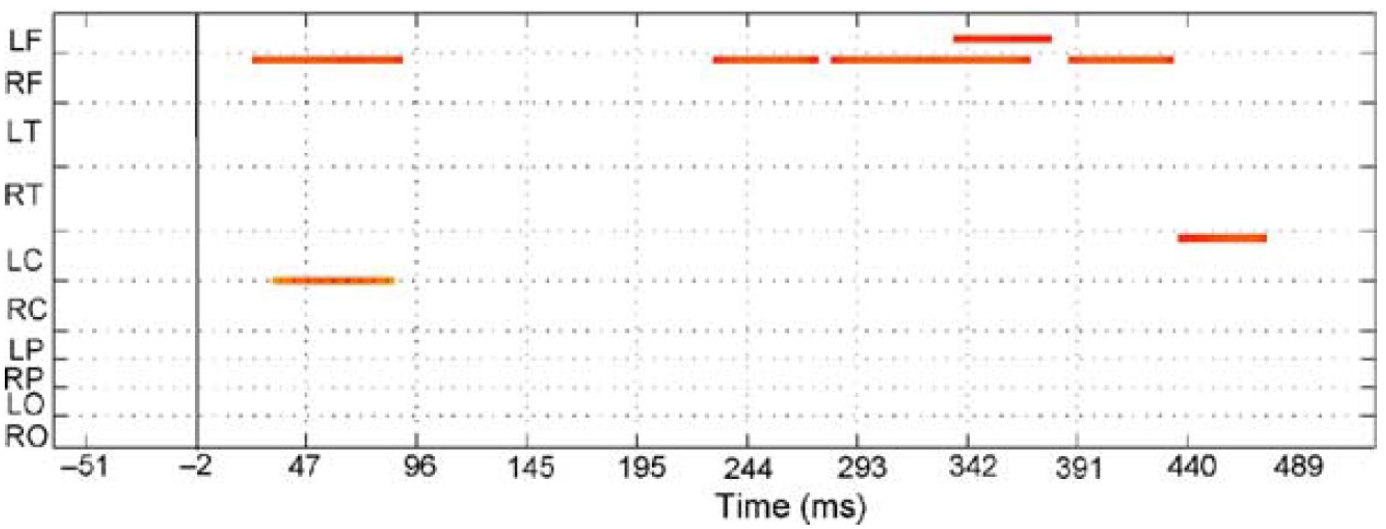
Significant crossmodal interactions displaying p-values for AV-(A+V), for the case when no illusion was perceived. Reproduced, with permission, from [3]. When the illusion is not perceived, only a couple of regions show cross-modal activity.

